# A single-cell atlas of transcribed *cis*-regulatory elements in the human genome

**DOI:** 10.1101/2023.11.13.566791

**Authors:** Jonathan Moody, Tsukasa Kouno, Miki Kojima, Ikuko Koya, Julio Leon, Akari Suzuki, Akira Hasegawa, Taishin Akiyama, Nobuko Akiyama, Masayuki Amagai, Jen-Chien Chang, Ayano Fukushima-Nomura, Mika Handa, Kazunori Hino, Mizuki Hino, Tomoko Hirata, Yuuki Imai, Kazunori Inoue, Hiroshi Kawasaki, Toshihiro Kimura, Tomofumi Kinoshita, Ken-ichiro Kubo, Yasuto Kunii, Fernando López-Redondo, Riichiro Manabe, Tomohiro Miyai, Satoru Morimoto, Atsuko Nagaoka, Jun Nakajima, Shohei Noma, Yasushi Okazaki, Kokoro Ozaki, Noritaka Saeki, Hiroshi Sakai, Kuniaki Seyama, Youtaro Shibayama, Tomohisa Sujino, Michihira Tagami, Hayato Takahashi, Masaki Takao, Masaru Takeshita, Tsuyoshi Takiuchi, Chikashi Terao, Chi Wai Yip, Satoshi Yoshinaga, Hideyuki Okano, Kazuhiko Yahamoto, Takeya Kasukawa, Yoshinari Ando, Piero Carninci, Jay W. Shin, Chung-Chau Hon

## Abstract

Transcribed cis-regulatory elements (tCREs), such as promoters and enhancers, are fundamental to modulate gene expression and define cell identity. The detailed mapping of tCREs at single-cell resolution is essential for understanding the regulatory mechanisms that govern cellular functions. Prior tCRE catalogs, limited by bulk analysis, have often overlooked cellular heterogeneity. We have constructed a tCRE atlas using single-cell 5’-RNA-seq, capturing over 340,000 single-cells from 23 human tissues and annotating more than 175,000 tCREs, substantially enhancing the scope and granularity of existing *cis*-regulatory element annotations in the human genome. This atlas unveils patterns of gene regulation, revealing connections between broadly expressed promoters and cell type-specific distal tCREs. Assessing trait heritability at single-cell resolution with a novel tCRE module-based approach, we uncovered the nuanced trait-gene regulatory relationships across a continuum of cell populations, offering insights beyond traditional gene-level and bulk-sample analyses. Our study bridges the gap between gene regulation and trait heritability, underscoring the potential of single-cell analysis to elucidate the genetic foundations of complex traits. These insights set the stage for future research to investigate the impact of genetic variations on diseases at the individual level, advancing the understanding of cellular and molecular basis of trait heritability.

## Introduction

The human body comprises diverse and specialized cell types. Gene expression, which defines cell identity, is regulated by *cis*-regulatory elements (CREs), mostly promoters and enhancers. (Zhang *et al*., 2021; Ong and Corces, 2011). CREs control gene expression by recruiting transcription factors (TFs) and RNA polymerase II to initiate transcription of capped-RNA (Cho *et al*., 1997) at both promoters and enhancers (Andersson *et al*., 2014). Sequencing of RNAs 5’-end pinpoints transcriptional start sites (TSS) and thus transcribed CREs (tCREs). tCREs can be categorized based on their proximity to the annotated gene: proximal tCREs (P-tCREs), such as promoters, are close to the gene TSS, while distal tCREs (D-tCREs), like enhancers, are located further away. Previous studies using TSS profiling in bulk samples, notably CAGE (Murata *et al*., 2014), concentrated on tissue samples and a limited number of primary cell types, yielding cell population-averaged information and a restricted scope (Forrest *et al*., 2014 FANTOM5). Existing single-cell atlases, largely based on gene expression, lack alternative promoters and distal CREs (Eraslan *et al*., 2022; Domínguez Conde *et al*., 2022; THE TABULA SAPIENS CONSORTIUM, 2022; Suo *et al*., 2022) limiting our ability to decode the regulatory roles of CREs in defining cell type identity. Genome-wide association studies (GWAS) identified variants associated with traits and diseases (Liu *et al*., 2019) that are highly enriched in CREs. Chromatin accessibility assays are routinely employed to identify accessible CRE (aCRE) (Buenrostro *et al*., 2015). Despite this, a significant number of distal aCREs lack the epigenomic marks of active enhancers (Thibodeau *et al*., 2018). Although some of these elements may function as insulators (Kim *et al*., 2007) or silencers (Pang and Snyder, 2020), their overall relevance in gene regulation remains elusive, affecting their interpretability in trait-associated variants annotation.

Single-cell omics allows the quantification of transcriptome, epigenome, and chromatin interactions among individual cells (Buenrostro *et al*., 2015, Heumos *et al*., 2023; Gaulton *et al*., 2023). In particular, single-cell 5’ RNA-seq (sc-5’-RNA-seq) enables the concurrent detection and quantification of tCREs, alongside gene expression profiling in single cells (Kouno *et al*., 2019). In this study, we used sc-5’-RNA-seq to annotate 175,032 tCREs across 341,156 cells of 180 distinct cell types from 23 human tissues. Our analysis linked D-tCREs to their target promoters, revealing cell type-specific CRE usage patterns. We characterized tCRE modules and their associations to 63 different traits and diseases, highlighting their relevance in cell type-specific gene regulation and in disease predispositions. Based on tCRE module usage in single-cells, we introduced the novel ICE-CREAM score to assess trait heritability enrichment at the single-cell level, revealing nuanced trait-gene regulatory relationships across a continuum of cell populations. Moreover, by analyzing trait-associated variants within tCREs to unravel their regulatory impacts, we have deepened the understanding of how genetic associations contribute to disease at the molecular and cellular levels.

## Results

### Detection of tCREs using sc-5’-RNA-seq

Enhancer RNAs (eRNA) are generally thought to be non-polyadenylated (Andersson *et al*., 2014); therefore, we assessed the sensitivity of D-tCRE detection by sc-5’-RNA-seq, comparing oligo(dT) (sc-end5-dT) and random hexamer (sc-end5-rand) priming in human dermal fibroblasts (DBFM) and induced pluripotent stem cells (iPSC). Most signals were observed at gene TSSs for both protocols as expected (**Fig. 1a**). Both protocols detected P-and D-tCREs with a high degree of overlap (**Fig. 1b**) and strong correlation in expression levels (**Fig. 1c**). Moreover, both protocols recapitulated the bidirectional transcription of eRNAs defined by bulk-CAGE in a cell type-specific manner (**Fig. 1d**). The detection of eRNAs by sc-end5-dT is unexpected, and likely can be attributed to internal priming (La Manno *et al*., 2018; Gaidatzis *et al*., 2015). Notably, sc-end5-dT demonstrated greater sensitivity at the per-cell level, with similar read distribution profiles (Supplementary Fig. 1,2). These findings affirm the efficacy of sc-end5-dT in detecting both P-tCREs and D-tCREs, including eRNAs.

**Figure 1:**
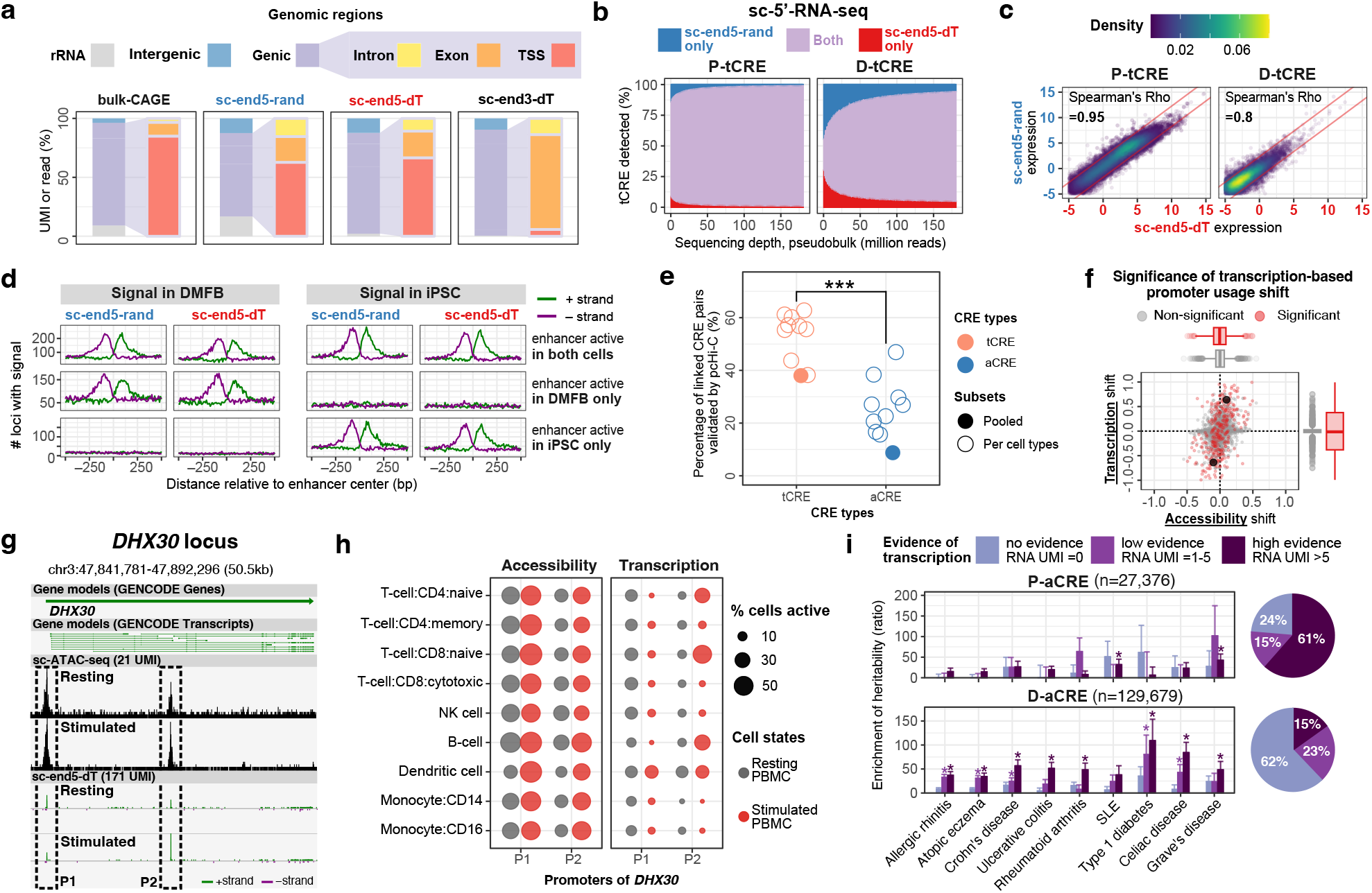
Detection of tCREs using sc-5’-RNA-seq. **a**) Distribution of reads aligning to the whole genome or to genic regions in bulk-CAGE and 5’-end random primed, 5’-end oligo(dT) primed and 3’-end oligo(dT) primed 10× single-cell RNA-seq. **b)** Proportion of overlap in tCRE detected in sc-end5-seq pseudo-bulk from 1 to 150 million reads. **c)** Correlation of tCRE levels between the pseudo-bulk data of the two sc-end5-seq methods. Red line, ±2-fold differences. UPM, UMI per million. **d)** TSS signal of sc-end5-dT and sc-end5-rand at bidirectionally transcribed enhancer loci defined in bulk-CAGE in iPSC and DMFB. **e)** Percentage of linked CRE pairs (co-activity score ≥0.2) validated (by pcHi-C) for tCRE (orange) and aCRE (blue), for per PBMC cell type (hollow circles) and for all cells pooled (solid circles). T-test for difference of tCRE and aCRE means shown. p <7×10^−6^, paired *t*-test for cell types. **f)** Shifts in alternative promoter usage upon stimulation for genes with multiple P-tCRE in CD8 T Cells. X-axis, change in accessibility (ratio of proportion of signal in sc-ATAC-seq) within tCRE upon stimulation; Y-axis, mean change in expression (ratio of proportion of signal in sc-end5-dT) of tCRE across meta-cells (k=50) upon stimulation. P, t-test for change in tCRE usage shown. Black dots highlight DHX30 promoters shown in g,h. **g)** Alternative promoter usage shift at DHX30 locus, modified from Zenbu genome browser view. **h)** Cell type-specific shift in alternative promoter usage at DHX30 locus. Proportion of cells with accessible aCRE (left) and transcribing tCRE (right) colored by stimulation state. **i)** Enrichment of heritability in aCREs with various levels of evidence of transcription. Y-axis, enrichment of heritability is measured as the ratio of proportion of heritability to proportion of SNP, estimated by LDSC. Error bars, standard error of the estimate. Asterisks, significant enrichments with p < 0.05.

We compared tCREs defined by sc-end5-dT with aCREs defined by sc-ATAC-seq in PBMCs under resting and activated states (Methods). Both methods offered similar cell clustering resolution, cell type specificity for CREs, and motif activity estimates (Supplementary Fig. 3). Using co-activity analysis (Pliner et al., 2018), tCRE pairs with high co-activity showed a greater validation rate via promoter-capture Hi-C (pcHi-C) (Javierre *et al*., 2016) (**Fig. 1e**). Upon PBMC activation, we identified 123 genes showing significant shifts in alternative promoter transcription, with only minimal changes in accessibility (**Fig. 1f**), as exemplified with the *DHX30* gene in CD8+ T-cells switching from promoter 1 to promoter 2 (**Fig. 1g-h**). This indicates that sc-ATAC-seq may have limited sensitivity in detecting changes in alternative promoter usage. Additionally, we found that increased transcriptional activity at aCREs correlated with enhanced trait heritability enrichment, particularly in distal aCREs (**Fig. 1i**). These findings highlight the capability of sc-end5-dT to capture cell type-specific P-and D-tCRE activities, leading to the creation of a comprehensive tCRE atlas using this approach.

### Annotating cell type clusters across 23 human tissues

We obtained sc-end5-dT single-cell or single-nuclei data, hereafter referred to as ‘single-cell’ data, from diverse human tissues via Single Cell Medical Network in Japan and public data (He, S *et al*., 2020) (Supplementary Table 1). Employing a standardized data processing pipeline for dataset integration (Methods), we constructed an atlas of 341,156 single-cells from 23 tissues (**Fig. 2a**). This atlas includes cells categorized into 21 Level 1 (Lv1) cell types (**Fig. 2b-c**, Supplementary Fig. 4) and further sub-clustering yielded 180 Level 2 (Lv2) cell types. To address sparsity and computational load while preserving transcriptional diversity, we created 3,350 meta-cells (**Fig. 2b**) (Supplementary Fig. 5,6). Analyses in this study were predominantly performed at the meta-cell level, unless specified otherwise.

**Figure 2:**
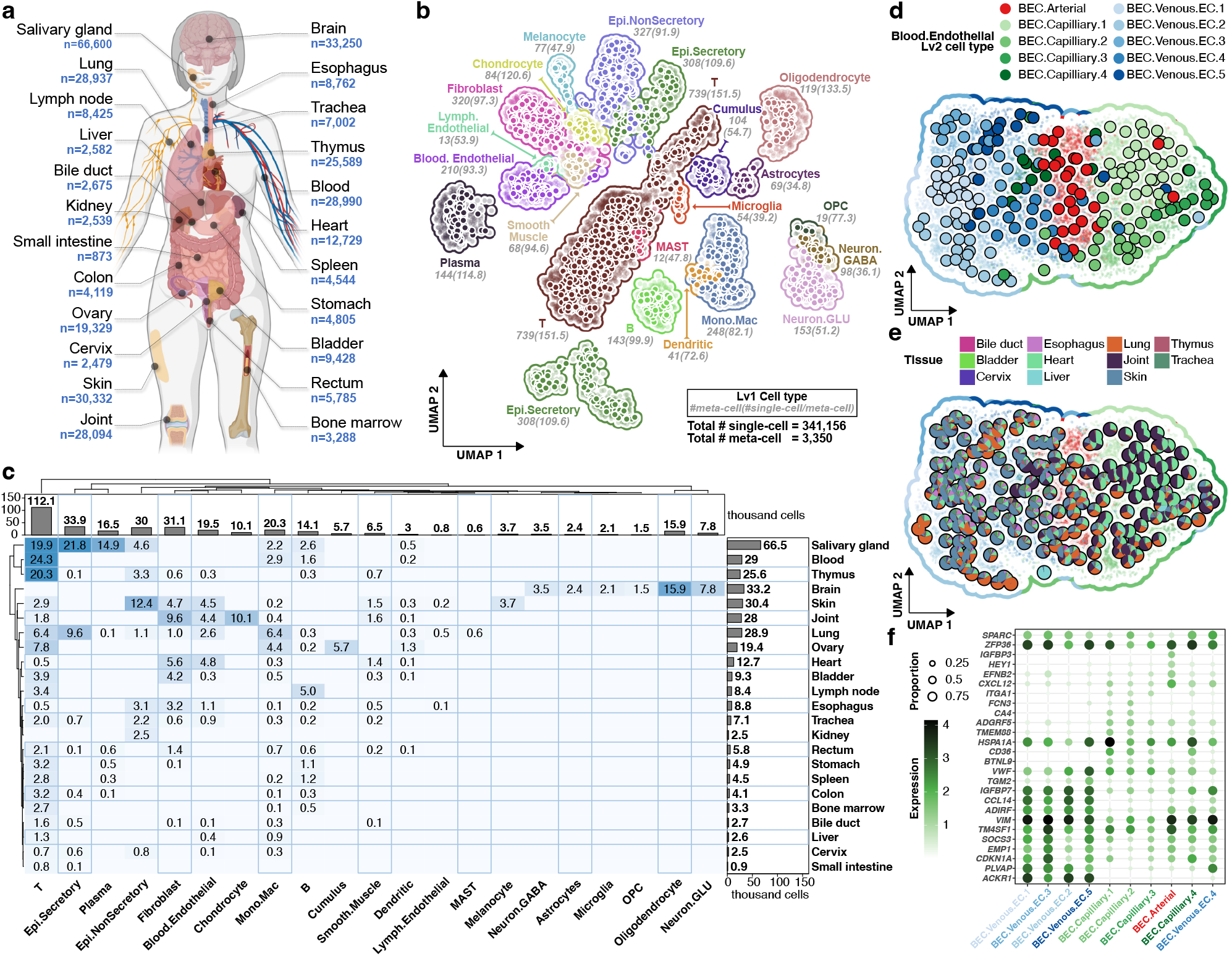
Annotating cell type clusters across 23 human tissues. **a**) Schematic with tissues of origin and number of included cells. **b)** Single-cell (small points) and meta-cell (large points) UMAP colored by Lv1 cell type clustering, meta-cells are positioned by the average UMAP positions of their single-cells, #meta-cells and average cells per meta-cell shown for each Lv1 cluster. **c)** Tissue of origin (rows) for cells in each Lv1 cell type (columns, in thousands of cells). **d,e)** Lv1 BEC subset reclustered and colored by Lv2 cell type cluster (d) and tissue of origin (e). **f)** Dotplot displaying top differentially expressed genes for each BEC Lv2 cluster.

To illustrate our cell annotations, we highlighted blood endothelial cells (BECs), distinguishing arterial, capillary, and venous subtypes, their tissue distribution, and marker genes in Level 2 (Lv2) cell types (**Fig. 2d-f**). For example, general capillary BECs displayed gene expression profiles indicative of inflammatory response and lipid transcytosis, marked by genes such as *BTNL9*, *ITGA1*, and *CD36*. Lung-enriched BEC.Capillary.2 subtypes were characterized by the pulmonary marker *CA4*. Notably, we observed an enrichment of capillaries in the heart and joint (BEC.Capillary.1) whereas venous BECs were enriched in the skin (BEC.Venous.1 and BEC.Venous.4) (**Fig. 2e**) (He, Y *et al*., 2022), aligning with the role capillary-to-myofiber interface plays in muscle function (Lemieux and Birot, 2021). Additionally, venous BECs showed higher expression of *CD74*, *CCL14*, *ACKR1* compared to arterial and capillary subtypes, suggesting a role in immune cell migration (Li *et al*., 2022). Detailed markers and tissue composition maps for Lv2 cell types highlight the diversity captured across immune, neuronal, stromal and endothelial cell types (Supplementary Fig. 6,7). In summary, these results demonstrated the utility and relevance of our cell type clustering and annotations.

### Building a single-cell tCRE atlas

Utilizing our single-cell data, we built a tCRE atlas comprising 81,829 proximal (P-tCREs) and 96,400 distal (D-tCREs) elements (Methods; Moody et al., 2022; Supplementary Table 2). The majority of these tCREs—94.3% of P-tCREs and 88.2% of D-tCREs were supported by candidate CREs from external epigenomic datasets from ENCODE (ENCODE Project Consortium *et al*., 2020) and a sc-ATAC atlas (Zhang *et al*., 2021), affirming the validity of our tCREs (**Fig. 3a**). The remaining unsupported tCREs may represent novel, cell type-specific elements. Notably, only 84.3% of P-tCREs and 46.7% of D-tCREs aligned with FANTOM5 TSS clusters (Forrest *et al*., 2014), expanding tCRE annotations within the human genome. Our analysis of cell type-specificity revealed a median enrichment of 7.8% for P-tCREs and 11.1% for D-tCREs in Lv1 cell types (**Fig. 3b**), with glutamatergic neurons displaying the highest specificity, consistent with known chromatin accessibility patterns (Hauberg *et al*., 2020), indicative of a relatively more complex gene regulatory architecture in glutamatergic neurons. Additionally, we categorized 66.1% of P-tCREs as gene promoters and the remainder as ‘flanking’, identifying 8,791 potential novel alternative promoters not listed in GENCODEv32. Overall, our atlas provides promoter annotations for 31,594 genes, including 12,386 with multiple promoters, averaging 4.4 promoters per gene (Supplementary Table 3).

**Figure 3:**
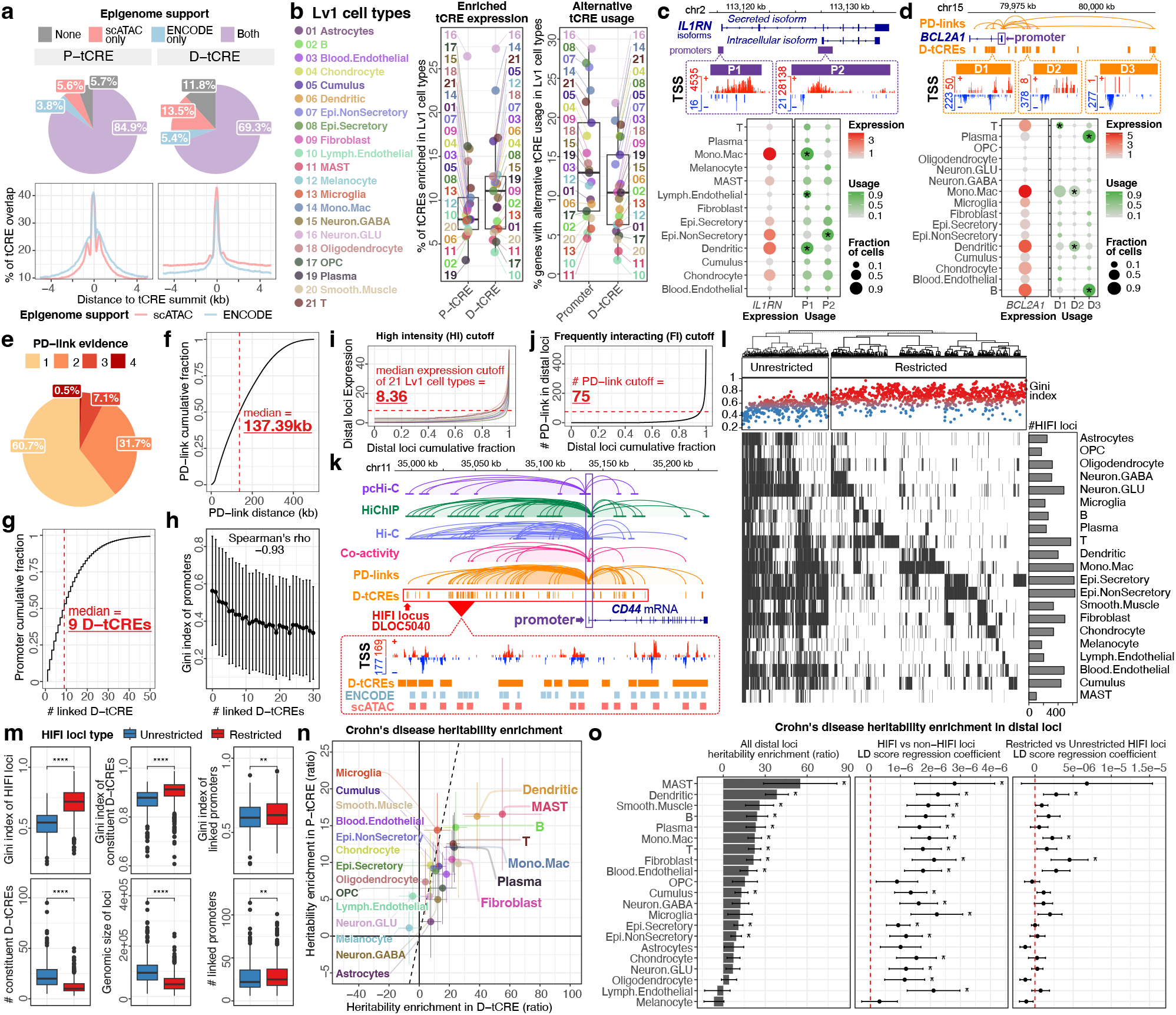
Building a single-cell tCRE atlas. **a**) Epigenome support of the tCREs. Percentages (upper) and coverage pattern (lower) of P-tCREs and D-tCREs overlap with ENCODE and sc-ATAC CREs. **b**) Cell type-specific expression of tCREs. Left, percentage of active P-tCREs and D-tCREs significantly enriched in Lv1 cell types; Right, percentage of genes with significant alternative usage of promoters and linked D-tCREs in Lv1 cell types. (p < 0.05, Wilcoxon test) **c**) Alternative promoter usage by *IL1RN.* An asterisk represents significant alternative promoter usage (p < 0.05, Wilcoxon test). **d**) Alternative D-tCRE at *BCL2A1*. An asterisk represents significant alternative D-tCRE usage (p < 0.05, Wilcoxon test). **e**) Corroboration of PD-links by pcHi-C, HiChIP, Hi-C, and Co-activity. **f**) Genomic distance of PD-links. **g**) Number of D-tCRE linked per promoter. **h**) Number of D-tCRE linked to promoters stratified by promoter Gini index. **i)** Expression cutoffs for high intensity distal loci, lines for each Lv1 cell type. Red dotted line, median of the cutoffs. **j)** Number of PD-link cutoffs for frequently interacting distal loci. **k)** HIFI locus at *CD44* region. **l)** Expression of HIFI loci. Right, the number of active HIFI loci in each Lv1 cell type. Heatmap k-mean clustered with k=2. Top, the Gini index of each HIFI locus from expression across Lv2 cell types. **m)** Comparisons between cell type-unrestricted and –restricted HIFI loci. Wilcoxon test. **n)** CD heritability enrichment in P-tCREs and D-tCREs. tCREs active in each Lv1 cell type were used to estimate heritability enrichment. **o)** CD heritability at distal loci. Left, heritability enrichment of all distal loci active in each Lv1 cell type. Middle, LD score regression coefficient comparing HIFI loci against non−HIFI loci. Right, LD score regression coefficient comparing restricted HIFI loci against unrestricted HIFI loci. Dots and error bars, estimated values and standard errors. An asterisk represents p < 0.05 in all cases. Selected transcripts shown. All boxes represent 25th, 50th and 75th percentile of the data.

Alternative promoter usage is a key mechanism for expanding transcriptome diversity and generating functionally distinct isoforms (Singer *et al*., 2008). On average 12.9% of multi-promoter genes (n=1,948 in total) exhibited significant alternative promoter usage across Lv1 cell types (**Fig. 3b**; Supplementary Table 4). The *IL1RN* gene, for example, employs distinct promoters for its secreted (P1) and intracellular (P2) isoforms, with P1 enriched in immune cells and P2 in non-secretory epithelial cells (**Fig. 3c**), indicating cell type-specific functionalities (Butcher *et al*., 1994) and aligning with the hypothesis that the intracellular form modulates IL-1 production in keratinocytes (Arend and Guthridge, 2000). Additionally, TF binding motif (TFBM) activity estimations suggested that differential promoter usage may be influenced by cell type-specific TF activity, with 48.5% (n=944 of 1,948) of genes with alternative promoters having significantly upregulated TFBMs in corresponding Lv1 cell types (Supplementary Fig. 8), indicating a TF-driven mechanism underpinning cell type-specific promoter usage.

We integrated three public chromatin interaction datasets with co-activity data from our atlas to infer Promoter-to-Distal tCRE interactions (PD-links), cataloging 466,079 PD-links for 75% of promoters (n=40,626) (Supplementary Table 5). Notably, 39% of these links were supported by at least two out of four evidence lines (**Fig. 3e**), with promoters connecting to a median of nine D-tCREs at a distance of 137.39 kb (**Fig. 3f-g**). Alternative usage of distal regulatory elements has broad implications for cell type identity, differentiation, and development (Nord *et al*., 2013). Our findings suggest that promoters with broader expression profiles across Lv2 cell types, indicated by a lower Gini index, are linked to more D-tCREs (**Fig. 3h**), suggesting extensive use of distal elements for regulating genes with a broad cellular activity. Furthermore, 10.4% of genes with multiple D-tCRE links showed significant changes in D-tCRE usage across Lv1 cell types (**Fig. 3b**). For example, the *BCL2A1* gene, pivotal for T cell development and survival (Mandal *et al*., 2005), exhibited differential D-tCRE usage correlating with its enriched expression pattern across immune cells (**Fig. 3d**). These results highlight that *BCL2A1* consistently maintains enriched expression across immune cell types, while it harbors unique sets of distal regulatory elements within each cell type, reinforcing the observation in Fig. 3h that the cell type-specific gene regulation is supported by distinct sets of D-tCRE.

In our atlas, we observed regions with intense D-tCRE activity and high frequencies of chromatin interactions, termed High Intensity and Frequently Interacting (HIFI) loci, (**Fig. 3k**; Supplementary Table 6), analogous to super-enhancers and FIREs (Schmitt *et al*., 2016; Hnisz *et al*., 2013). For example, the *CD44* region contains a HIFI locus (DLOC5040) with 47 D-tCREs spanning 186.6 kb. Most of these D-tCREs display bidirectional transcription and are supported by epigenomic data, with 74.4% (35 of 47) linked to the *CD44* promoter, as corroborated by coactivity and chromatin interaction data (**Fig. 3k**). We cataloged 1,229 HIFI loci, with each Lv1 cell type expressing a median of 336 HIFI loci (**Fig. 3l**). These were classified as either cell type-unrestricted (n=377) or –restricted (n=852) based on their expression patterns, correlating well with Gini index distributions (**Fig. 3l-m**). At unrestricted loci, both D-tCREs and their linked promoters showed significantly lower Gini indices compared to restricted loci (**Fig. 3m**), suggesting a role for distal elements in gene expression refinement and specificity across cell types. The unrestricted loci also comprise more D-tCREs and span larger genomic regions, implying a more complex regulatory mechanism at these loci across cell types (**Fig. 3m**).

To assess the biological relevance of various tCRE categories, we investigated their enrichment in trait and disease heritability (Finucane *et al*., 2015). We observed similar enrichment levels for both P-and D-tCREs across Lv1 cell types (Supplementary Fig. 9). In immune cells, tCREs exhibited higher enrichment in Crohn’s disease (CD) heritability, particularly D-tCREs (e.g., dendritic cells in **Fig. 3n**), which is consistent with their critical role in microbial recognition and innate immunity (Bates and Diehl, 2014). Additionally, cell-type-specific trait enrichments, such as in BECs and smooth muscle cells (SMCs) for varicose veins, and microglia and oligodendrocyte progenitors for Parkinson’s disease (PaD), were observed (Supplementary Fig. 9). CD heritability enrichment was notably higher at HIFI loci compared to non-HIFI loci (**Fig. 3o**), mirroring the enriched disease heritability observed in super-enhancers (Hnisz *et al*., 2013). Further, cell type-restricted HIFI loci were more enriched in heritability within relevant cell types, like dendritic cells, monocytes/macrophages, and fibroblasts, highlighting the cell type-specific importance of these loci (**Fig. 3o**; Supplementary Fig. 10 for all other traits). These findings underscore the critical role of distal regulatory elements in the cell type-specific landscape of disease heritability.

### Inferring regulatory programs with tCRE modules

Applying consensus Non-negative Matrix Factorization (cNMF) to our meta-cell data, we identified 150 tCRE regulatory modules that represent independent biological properties in specific cell populations, such as muscle contraction in SMCs (Kotliar *et al*., 2019) **(Fig. 4a,f)**. These modules are largely cell type-specific (Supplementary Fig. 11), with, for example, M011 being specific to BEC subsets, while M033 is specific to fibroblasts (**Fig. 4b-e**). Further analysis within the stromal cell subset, including SMCs, lymph endothelial cells (LECs), and chondrocytes, pinpointed modules like M053 and M028 as SMC-specific, related to muscle function and cardiac biology, and notably enriched in myocardial infarction (MI) heritability (**Fig. 4f**), underscoring the protective role of SMCs in mediating superoxide free radicals within the aortic wall (Zhuge *et al*., 2020). Additionally, significant MI heritability enrichment was observed in one BEC-associated module (M011) and two fibroblast-associated modules (M012 and M080). These findings provide insights into tCRE module usages within SMCs, BECs, and fibroblasts and suggest their relevance to MI, underscoring the biological significance of the tCRE modules we identified. Moreover, our analysis delineates tissue-specific module-trait relationships across immune, neuronal, and epithelial cells (Supplementary Fig. 12-14), reinforcing the intricate cell type-specific nature and disease relevance of these tCRE modules.

**Figure 4:**
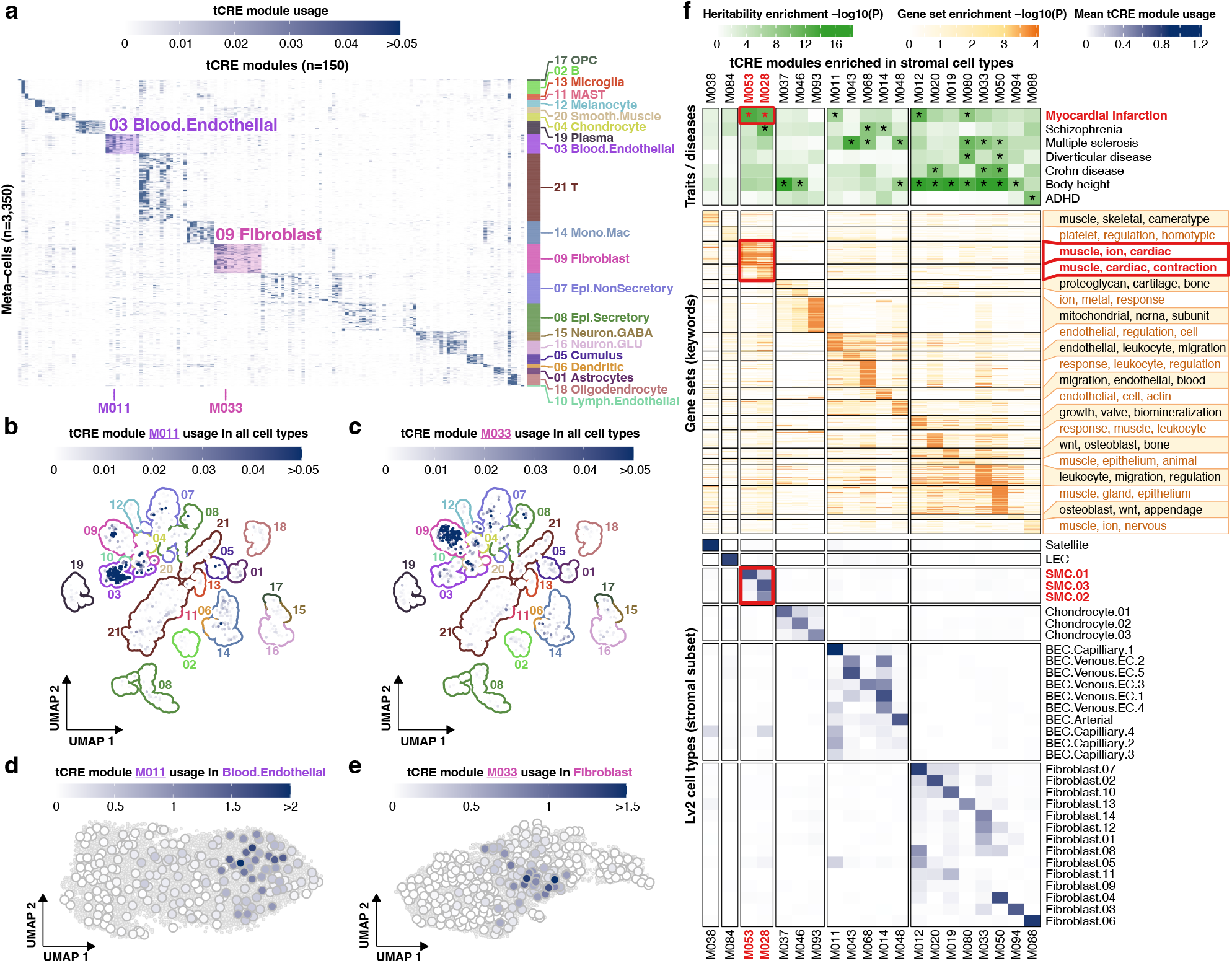
Inferring regulatory programs with tCRE modules. **a**) Heatmap of module usage in 150 tCRE modules (columns) in meta-cells (rows) annotated (right) with Lv1 cell type clusters. **b)** UMAP plot of module M011 usage. **c)** UMAP plot of module M033 usage. **d)** UMAP plot of M011 usage within the Lv1 BEC cluster. **e)** UMAP plot of M033 usage within Lv1 fibroblast cluster **f)** tCRE modules enriched in stromal cells (columns across 3 heatmaps) with: *i)* Heritability enrichment: for selected traits with significant enrichment in the stromal cell subset. *ii)* Gene set enrichment: –log10(FDR) of GO terms amongst P-tCREs, over-represented keywords are shown. *iii)* Module usage: mean meta-cell module usage in Lv2 cell type clusters.

### Assessing trait heritability at single-cell resolution using ICE-CREAM score

Identifying cell types implicated in diseases is crucial for biomedical research. We have developed an analytical framework to assess trait heritability enrichment at the single-cell or meta-cell level based on trait heritability enrichment in tCRE modules. This allows for interrogation of trait heritability in a manner dependent or independent of cell type annotations. In this framework, we calculate a trait heritability enrichment score, the ICE-CREAM score, for each cell by summing the usage of all modules weighted by their heritability enrichment for a trait, then evaluating the significance against a permuted null distribution, with score expressed as –log10(p-value) (Methods). Applying the ICE-CREAM score to analyze 63 traits across 3,350 meta-cells revealed the specificity of cell types to these traits (**Fig. 5a**). When projected onto single cells, similar patterns were observed (Data availability). Using Cell-Set Enrichment Analysis (CSEA) to quantify trait enrichment in Lv1 cell types (Supplementary Table 8), we identified a link between COVID-19 severity and monocyte/macrophage cells, consistent with their documented recruitment in severe cases (Zhou *et al*., 2020), and with BECs and mast cells, known to be implicated in COVID-related thrombosis (Afrin *et al*., 2020; Bonaventura *et al*., 2021). Moreover, our approach revealed the involvement of diverse cell types in complex diseases, as evidenced by the enrichment across immune cells, fibroblasts, SMCs, and endothelial cells in psoriasis (**Fig. 5a**).

**Figure 5:**
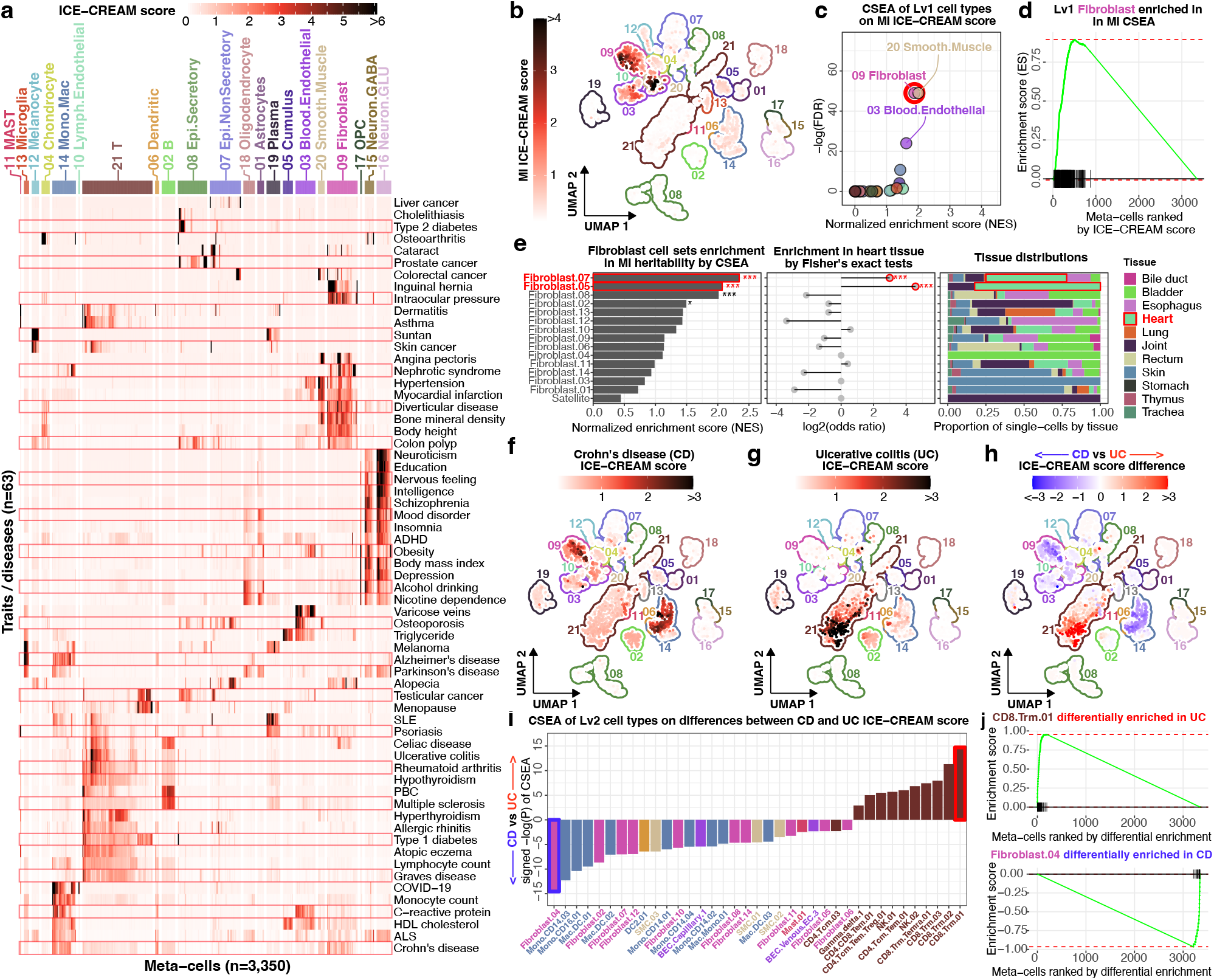
Assessing trait heritability at single-cell resolution using ICE-CREAM score. **a**) Heatmap of ICE-CREAM score for traits (rows) in meta-cells (columns) colored by Lv1 cell type clusters (above). **b)** UMAP of MI ICE-CREAM score in meta-cells. **c)** Lv1 CSEA for MI, x-axis NES, y-axis –log10(FDR) **d)** Ranking fibroblast meta-cells for MI enrichment in Lv1 CSEA. **e)** (left) Lv2 fibroblast cell type enrichment for MI in CSEA, (center) Enrichment for Heart cells, an asterisk represents significant enrichment (p < 0.05, Fisher’s exact test) (right) tissue of origin proportion. **f)** UMAP of CD ICE-CREAM score. **g)** UMAP of UC ICE-CREAM score. **h)** UMAP of difference between CD and UC ICE-CREAM score. **i)** Lv2 cell type clusters with significant difference in ICE-CREAM scores. **j)** CSEA of meta-cells corresponding to the most divergent Lv2 cell type clusters in (h,i).

Highlighting MI, we noted significant heritability enrichments within Lv1 cell types of BECs, fibroblasts, and SMCs (**Fig. 5b-d**), aligning with the module enrichments depicted in Fig. 4f. Further CSEA of Lv2 fibroblast cell types pinpointed MI heritability enrichment particularly in Fibroblast.07 and Fibroblast.05, which were notably prevalent and significantly enriched in heart tissues (**Fig. 5e**). These findings illustrate the role of tissue origin and microenvironment in influencing cell subtype specification and their contributions to disease. For a more detailed understanding of cell type-specific trait heritability, we extended the CSEA to Lv2 cell types for all traits studied, offering a high-resolution view of cell type to trait associations (Supplementary Table 9; Supplementary Fig. 15-18).

A pairwise comparison of ICE-CREAM scores for closely related traits elucidated fine-grained differences in cell type relevance between diseases. For example, when contrasting CD and ulcerative colitis (UC), two related immune disorders affecting different parts of gastrointestinal tract, we found CD heritability to be more enriched in monocyte, macrophage, and fibroblast subtypes (**Fig. 5f**), correlating with the significant role of monocytes in CD (Grip *et al*., 2007) and fibrogenesis in CD-associated intestinal fibrosis (Burke *et al*., 2007). In contrast, UC showed higher heritability enrichment in CD8+ memory and NK cells, underscoring the contribution of NKT cells to the atypical TH2 response in UC (Fuss *et al*., 2004) (**Fig. 5g**). We quantified these differences by applying CSEA to differential ICE-CREAM score rankings between CD and UC, which highlighted CD8.Trm.01 and Fibroblast.04 as the most differentially enriched Lv2 cell types for CD and UC, respectively (**Fig. 5h-j**). Notably, module M050, which is highly specific to Fibroblast.04 and enriched in CD heritability, showed enrichment in epithelium-related gene sets (**Fig. 4f**), aligning with the proposed involvement of epithelial fibroblasts in CD (Burke *et al*., 2007).

Neurological traits such as schizophrenia, insomnia, and neuroticism showed strong associations with GABAergic and glutamatergic neurons, while neurodegenerative diseases like Alzheimer’s diseases and PaD correlated with microglial activity (**Fig. 5a**). In contrasting amyotrophic lateral sclerosis (ALS) with PaD, PaD was notably enriched in oligodendrocytes and microglia, aligning with evidence of microglial activation and consequent neuronal damage in PaD (Bae *et al*., 2023; Long-Smith *et al*., 2009; Hickman *et al*., 2018; Muzio *et al*., 2021), while ALS showed enrichment in dendritic cells and macrophages, known for their inflammatory role in ALS (Rusconi *et al*., 2017) (Supplementary Fig. 19). Additionally, in contrasting hypertension with varicose veins, the latter showed greater enrichment in subsets of BECs, whereas hypertension was more associated with fibroblasts and SMCs, which is consistent with their roles in vascular function (Touyz *et al*., 2018) (Supplementary Fig. 20). Overall, these results highlight the value of the ICE-CREAM score in identifying specific cell types contributing to traits, advancing our understanding of disease mechanisms at the cellular level.

### Linking trait-associated variants to relevant cell populations, genes and CREs

To elucidate genetic associations with traits, we prioritized trait-associated variants residing in tCREs using ICE-CREAM scores, genomic context, PD-links, and TFBM activity (Methods). We specifically examined SNPs that disrupt TFBMs in relevant cell types and those within HIFI loci, which exhibit high heritability enrichment (**Fig. 3o**). Approximately 66% of trait-associated loci (median per trait) were annotated with at least one SNP in a tCRE enriched in relevant cell types, as determined by ICE-CREAM score CSEA. In addition, ∼56% of trait-associated loci contained at least one SNP disrupting a TFBM correlated with trait heritability (Supplementary Fig. 21, Supplementary Table 10, 11).

To illustrate the value of these annotations, consider rheumatoid arthritis (RA), where T cells were identified as the most strongly associated Lv1 cell type (**Fig. 6a**). At RA risk loci, we focused on HIFI loci, particularly DLOC24008 near *PTGER4*, which showed a high correlation with the RA ICE-CREAM score and specificity to T cells (**Fig. 6b,c,e**), in contrast to the broadly expressed *PTGER4* (**Fig. 6d,f**). Interestingly, a large fraction of D-tCREs within DLOC24008 linked to the *PTGER4* promoter (**Fig. 6h**), underscoring how genes with broad expression patterns can achieve cell type-specific regulation through distal tCREs. The RA-associated SNP rs6883964 disrupts an *IRF1* motif within DLOC24008 (**Fig. 6h**) and the *IRF1* motif is highly active in immune cells (**Fig. 6g**). The documented associations of genomic region to multiple immune traits (Libioulle *et al*., 2007; Rodriguez-Rodriguez *et al*., 2015) substantiates the functional association of this SNP with *PTGER4* expression specifically in immune cells. This case demonstrates how D-tCREs confer cell type-specific regulation to broadly expressed genes and aids in the interpretation of non-coding SNPs in intergenic regions with cellular contexts.

**Figure 6:**
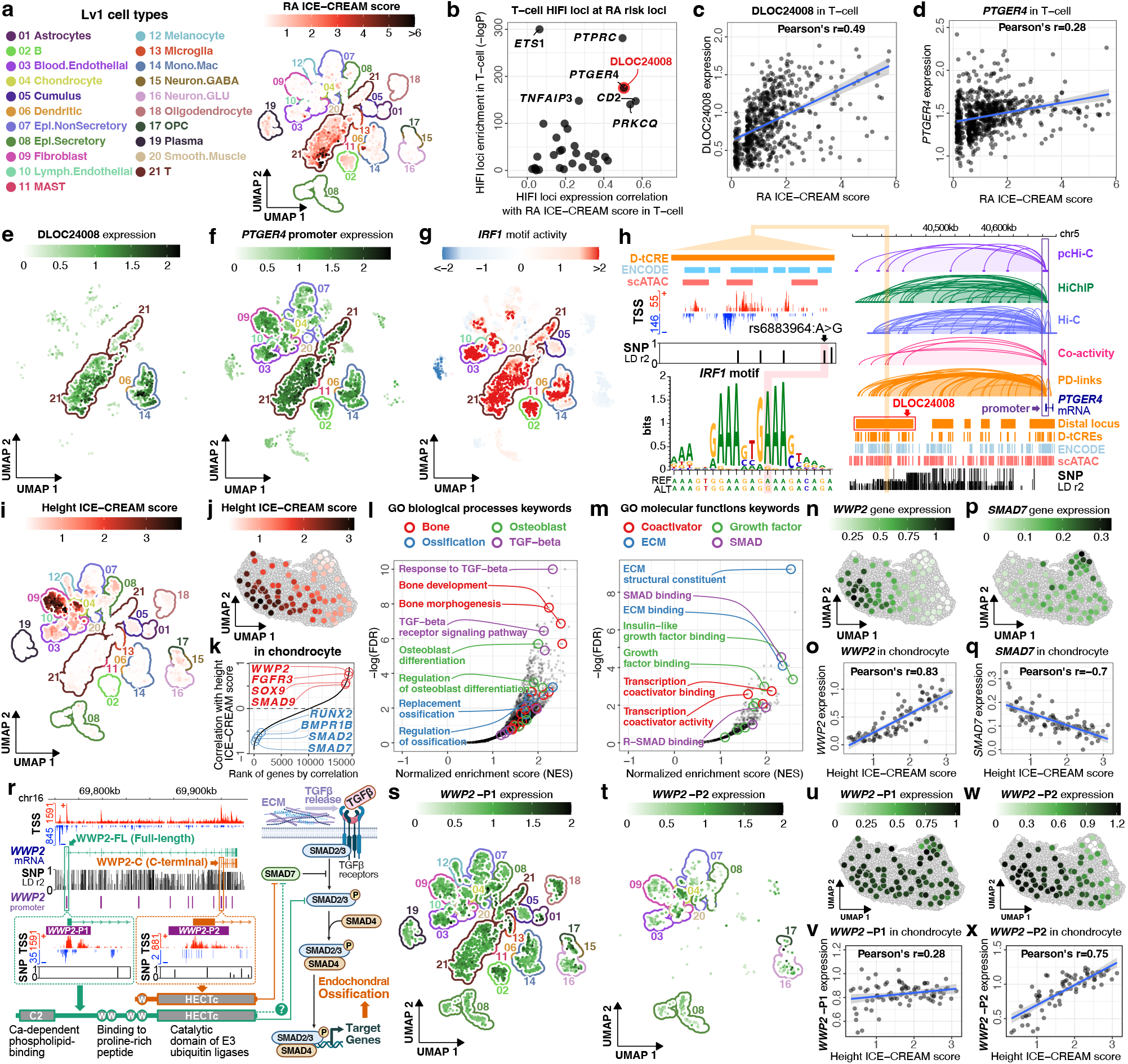
Linking trait-associated variants to relevant cell populations, genes and CREs. **a**) RA ICE-CREAM score UMAP, **b)** HIFI loci scatter plot on x-axis correlation with RA ICE-CREAM score, y-axis enrichment within T-cells (Wilcoxon rank-sum test). **c)** HIFI locus DLOC24008 Pearson’s correlation with RA ICE-CREAM score in meta-cells. **d)** *PTGER4* promoter Pearson’s correlation with RA ICE-CREAM score in meta-cells. **e)** HIFI locus DLOC24008 expression UMAP, summed expression of contained D-tCRE. **f)** *PTGER4* promoter expression UMAP. **g)** IRF1 motif activity UMAP. **h)** *PTGER4* gene and DLOC24008 genome browser view. **i)** Height ICE-CREAM score UMAP. **j)** Height ICE-CREAM score in chondrocyte Lv1 cluster. **k)** Ranked gene expression Pearson’s correlation with height ICE-CREAM score in chondrocyte meta-cells. **l,m)** GSEA enrichment for GO biological processes (l) and GO molecular functions (m) ranked by abs(gene expression) correlation with height ICE-CREAM score in chondrocyte Lv1 cluster. **n)** WWP2 gene expression in chondrocyte Lv1 cluster. **o)** Pearson’s correlation of *WWP2* and height ICE-CREAM score in chondrocyte Lv1 cluster. **p)** Ranked gene expression Pearson’s correlation with height ICE-CREAM score in chondrocyte meta-cells. **l)** *SMAD7* gene expression in chondrocyte Lv1 cluster. **q)** Pearson’s correlation of *SMAD7* and height ICE-CREAM score in chondrocyte Lv1 cluster. **r)** *WWP2* genome browser view (left) with TSS signal, highlighting P1 and P2 producing the WWP2-FL and WWP2-C isoforms respectively. (right) Schematic of WWP2 regulation of SMAD degradation in TGF-β signaling. **s,t)** *WWP2* P1 (s) and P2 (t) expression UMAP. **u,w)** *WWP1* P1 (u) and P2 (w) expression in the chondrocyte Lv1 cluster. **v,x)** *WWP1* P1 (v) and P2 (x) Pearson’s correlation with ICE-CREAM score in chondrocyte Lv1 cluster meta-cells.

We probed trait-associated SNPs in alternative promoters, uncovering significant heritability enrichment for body height trait within fibroblast and chondrocyte Lv1 cell types (**Fig. 6i**), highlighting the crucial role of chondrocytes in endochondral ossification: a process by which growing cartilage is systematically replaced by bone to form the growing skeleton. Chondrocyte meta-cells displayed a gradient of height ICE-CREAM scores that correlate with the expression of essential ossification regulators *SOX9* (Hattori *et al*., 2010) and *RUNX2* (Chen *et al*., 2014) (**Fig. 6j-k**). A GSEA, where we ranked the absolute correlation between gene expression and height ICE-CREAM score, further underscored the involvement of biological processes and molecular functions tied to bone biology and the critical components of the TGF-β signaling pathway (**Fig. 6l-m**), e.g. SMAD and extracellular matrix (**Fig. 6r**) (Estrada *et al*., 2013; Mokuda *et al*., 2019).

The inverse expression patterns and trait correlations between *WWP2* and *SMAD7* in chondrocyte meta-cells underscore the ubiquitination of SMAD7 by WWP2 within the TGF-β pathway (**Fig. 6n-r**). Two promoters lead to different WWP2 isoforms (de Kroon *et al*., 2017; Soond and Chantry, 2011; Wahl *et al*., 2019): Promoter 1 (P1) produces a full-length isoform (WWP2-FL) with broad expression, whereas Promoter 2 (P2) generates a chondrocyte-enriched shorter isoform (WWP2-C), with P2 expression strongly correlated with the height ICE-CREAM score, but not P1 (**Fig. 6r-x**). The observed gradient in height ICE-CREAM score may be influenced by the selective binding of SMAD proteins to the WWP2 isoforms, particularly the affinity of WWP2-C for SMAD7, impacting TGF-β signaling in endochondral ossification and ultimately skeletal growth and body height (de Kroon *et al*., 2017; Wahl *et al*., 2019) (**Fig. 6r**). These detailed tCRE-based analyses provide a nuanced understanding of trait associations, offering insights beyond traditional gene-level analyses.

## Conclusions

This single-cell tCRE atlas marks a considerable advancement over our previous efforts on bulk sample (Forrest *et al*., 2014), expanding the scope to include a wider array of tCREs and cell types, and enhancing granularity to single-cell resolution. This substantially improved the depth and breadth of tCRE annotations within the human genome. By interrogating distal regulatory elements and their associated promoters, our analyses revealed underlying patterns of gene regulation, such as the connection between broadly expressed promoters and cell type-restricted D-tCREs (**Fig. 3h**). The integration of tCRE information into trait heritability assessments through the ICE-CREAM score reveals subtle trait associations across cell populations (e.g. body height heritability in chondrocytes, **Fig. 6j**; WWP2 promoter effects, **Fig. 6r**), offering fresh insights into gene regulation and trait heritability in continuous cell populations. While current approaches like sc-linker (Jagadeesh *et al*., 2022) and h-magma (Sey *et al*., 2020) analyze trait-associated SNPs within regulatory elements but overlook a continuum of cell populations, and scDRS (Zhang *et al*., 2022) considers the continuum but omits regulatory elements, our approach addresses both, providing higher resolution and functional interpretability in a more flexible framework. Although sc-ATAC-seq is a prevalent technique for studying CREs at single-cell resolution, our data indicate that most distal aCREs are not transcribed (**Fig. 1i**), whereas transcribed aCREs show a greater enrichment for trait heritability (**Fig. 1i**). The functional significance of untranscribed distal aCREs in gene regulation remains to be fully understood, yet our findings underscore the value of transcriptional signals in studying CREs, particularly concerning trait heritability. Looking forward, it is imperative to evaluate the applicability of our findings at the individual level by single-cell tCRE profiling on a population scale and to investigate how genetic variants influence CRE activities and disease predispositions in specific cell types for diagnostic and therapeutic advancements. In conclusion, our work highlights the power of sc-5’-RNA-seq in mapping tCREs across cell types and advancing our understanding of the genetic, molecular and cellular drivers of diseases and traits.

## Methods

### Human Subjects

All human samples examined in this study were either exempted material or were obtained with informed consent and covered under the following research protocols: RIKEN Yokohama Campus (no. H28-24, H30-9, H30-26), Ehime University Hospital (1812005), Keio University Hospital (20170302, 20160377), Keio University School of Medicine (2019-0212), The Jikei University School of Medicine (33-438(11065)), Osaka University Hospital (21113-2), the University of Tokyo (2018192G-(4)). Written informed consent on sample collection, data acquisition and usage, and publication was obtained from all the participants.

### Single-cell 3’ and 5’ RNA-seq

Freshly prepared iPSC and DMFB cells were loaded onto the ChromiumTM Controller (10× Genomics®) on different days. Cell number and viability were measured by CountessTM II Automated Cell Counter (Thermo Fisher®). Final cell density was adjusted to 1.0×106cells/ml with >95% viability. Both cells were targeting ∼5,000 cells per reaction. For sc-end3-dT libraries, we used ChromiumTM Single Cell 3′ Library kit v2 (10× Genomics®). Briefly, single-cell suspensions were mixed with the Single-cell Master Mix using Reverse transcription (RT) Primer (AAGCAGTGGTATCAACGCAGAGTACATr–GrGrG) and loaded together with 3′ gel beads and partitioning oil into a Single Cell A Chips according to the manufacturer’s instructions (10× Genomics®). For sc-end5-dT and sc-end5-rand libraries, we used ChromiumTM Single Cell 5′ Library kit v1.1 (10× Genomics®). Single-cell suspension was mixed with Single-cell Master Mix using oligo(dT) RT primer (AAGCAGTGGTATCAACGCAGAGTACGAGAC–T(30)–VN) or random hexamer RT primer (AAGCAGTGGTATCAACGCAGAGTACNNNNNN) and loaded together with 5′ gel beads and partitioning oil into a Single Cell A Chips according to the manufacturer’s instructions. RNAs within single-cells were uniquely barcoded and reverse transcribed within droplets. Both methods used VeritiTM Thermal Cycler (Applied Biosystems®) for RT reaction. After collecting cDNAs prepared from each method, they were amplified using cDNA primer mix from the kit, followed by the standard steps according to manufacturer’s instructions. For iPSC and DMFB, six libraries (i.e. 3 methods × 2 cell lines) were barcoded by different indexes from i7 sample index plate (10× Genomics®). The libraries were examined in BioanalyzerTM (Agilent®) for size profiles and quantified by KAPATM Library Quantification Kits (Kapa Biosystems®). All libraries were sequenced on HiSeqTM 2500 (Illumina®) as 75 bp paired-end reads.

### Single-cell ATAC-seq

Freshly prepared resting and stimulated PBMCs were subjected to sc-end5-dT (Single Cell 5′ Library kit v1.1) and sc-ATAC-seq (Single Cell ATAC kit v1.1) library construction on the same day using the ChromiumTM platform according to manufacturer’s instructions (10× Genomics®). About 5,000 cells/nuclei were targeted per reaction. sc-end5-dT and sc-ATAC-seq libraries were sequenced on HiSeqTM 2500 (Illumina®) as 75bp and 100bp paired-end reads respectively.

### PBMC stimulation

Human PBMCs were prepared from the whole blood of a male healthy donor with LeucosepTM (Greiner®). Isolated 2×106 PBMC cells were incubated with PMA/ionomycin (i.e. stimulated) (Cell Activation Cocktail with Brefeldin A, Biolegend®), or DMSO as control (i.e. resting), for six hours.

### Bulk CAGE, RNA-seq and ATAC-seq library construction and sequencing for DMFB and iPSC

Bulk CAGE libraries were generated by the nAnT-iCAGE (Murata *et al*., 2014) method as previously described and sequenced on HiSeqTM 2500 (Illumina®) as 50bp single-end reads. Bulk RNA-seq libraries were generated as previously described (Andersson *et al*., 2014) and sequenced on HiSeqTM 2500 (Illumina®) as 100bp paired-end reads. Bulk ATAC-seq was performed as previously described (Buenrostro *et al*., 2015) with slight modifications. Briefly, 2.5×104 cells/ml were used for library preparation. Due to the more resistant membrane properties of DMFB, 0.25% IGEPALTM CA-630 (Sigma-Aldrich®) were used for cell lysis. Transposase reaction was carried out as described in the protocol followed by 10 to 12 cycles of PCR amplification. Amplified DNA fragments were purified with MinEluteTM PCR Purification Kit (QIAGEN®) and size-selected with AMPureTM XP (Beckman Coulter®). All libraries were examined in BioanalyzerTM (Agilent®) for size profiles and quantified by KAPATM Library Quantification Kits (Kapa Biosystems®). Bulk ATAC-seq libraries were sequenced on HiSeqTM 2500 (Illumina®) as 50bp paired-end reads.

### Processing sc-end5-dT data for PBMC

Reads were aligned to hg19 with *Cell Ranger* and the gene-based expression matrixes were processed with *Seurat v3*. Briefly, cells were excluded with ≥4 median absolute deviation from the mean for number of features, UMI count, and percentage of mitochondrial UMI. Top 2,000 variable features were selected. Resting and stimulated PBMC samples were integrated with Canonical correlation analysis (CCA) implemented in *Seurat* using principal component (PC) 1 to 20 based on gene-based expression matrix. Bam files were processed with *SCAFE (v1.0.0)* to generate filtered CTSS bed files and *de novo* define tCRE. tCRE-based expression matrices from *SCAFE* were added to the *Seurat* object for downstream analysis. Cell annotation was performed by manually combining annotations from *scMatch* (Hou *et al*., 2019) *(version at 2020-10-10)* and known marker genes. cell type-specificity and stimulation-specificity of tCREs were calculated with *Seurat FindMarkers* function with min.pct=0, return.thresh=Inf, logfc.threshold=0, min.cells.group=0.

### Processing sc-ATAC-seq data for PBMC

Reads were aligned to hg19 with *Cell Ranger ATAC v1*.*2* (10× Genomics) and the data were processed with *SnapATAC* (Fang *et al*., 2020) *v1*.*0*.*0* using default parameters, selecting cells with ≥40% reads in ATAC peaks. Resting and stimulated cells were integrated with *Harmony v1*.*0* using PC 1 to 20. sc-ATAC-seq and sc-end5-dT were integrated using *SnapATAC FindTransferAnchors* and *TransferData* functions to transfer cell cluster annotations from the sc-end5-dT cells to the sc-ATAC-seq cells. sc-ATAC-seq peaks were defined per cell type using *SnapATAC runMACS*, then merged. These merged peaks were referred to as aCREs and these aCREs were annotated using *SCAFE*. Cell type-specificity and stimulation-specificity of aCREs were calculated with *SnapATAC findDAR*.

### Analysis of DMFB, IPSC and PBMC data in Figure 1, Supplementary Figures 1-3

Reads were aligned to hg19 with *Cell Ranger v3*.*1*.*0* (10× Genomics), and bam files were processed with *SCAFE* to generate filtered CTSS bed files and *de novo* define tCRE. Annotation counts were produced by intersecting CTSS bed files with GENCODE gene models. Metagene plots from overlapping CTSS bed files with exons binned with Bioconductor *equisplit* using *foverlaps*. Enrichment of genesets in sc-end5-dT versus sc-end5-rand was tested using *fgsea v1*.*16*.*0* with nperm=1000. Genesets were defined as: 1) cytoplasmic, nucleoplasmic, and chromatin-bound RNAs: log_2_ fold-change ≥2 in fractionated CAGE data compared to total CAGE data, 2) long and short RNAs: maximum transcript length per gene ≥25,000nt and <1,000nt, 3) Non-polyA histone RNAs: histone RNAs with log2 fold-change ≥2 in non-polyA fraction in a previous study (Yang et al., 2011) *ChromVAR v1*.*12*.*0* was used to estimate per-cell TF motif activities for the JASPAR2018 core motif set for tCRE or aCRE excluding chrM. The tCRE expression matrix was binarized prior to running. *Cicero v1*.*3*.*4*.*11*was used to calculate the co-activity score between CRE pairs using default parameters. Only tCREs and aCREs present in ≥3 cells were considered. Co-activity scores were estimated separately using cells within individual cell types (cell type sets) or all cells (pooled set). A pair of CREs with co-activity score ≥0.2 is defined as “linked”. pcHi-C connections (without cutoffs) from all cell types were pooled and used for validation of co-activity linked CREs pairs. For comparisons of validation rates between tCREs and aCREs, only a subset of CREs that are overlapped between tCREs and aCREs and CRE pairs located ≥10kb apart was used. Detecting shifts in alternative promoter use: For each cell type (excluding dendritic cells due to low cell count), knn clustering of the *Seurat* SNN matrix (k=50) was used to generate meta-cells. The proportion of UMI in each gene arising from P-tCREs was calculated for each meta-cell. cell type-specific tCRE switching events were identified using a *t*-test for differences in the proportion of UMI in gene contributed from each tCRE between meta-cells of selected cell type and a background of all other cell types. sc-ATAC-seq signal (UMI per millions) at a tCRE was defined as the maximum signal in cell type bigwig files generated with *SnapATAC runMACS*.

### tCRE atlas scRNA alignment, filtering, doublet removal, processing

Fastq from the Single Cell Medical Network in Japan were aligned to hg38 using cellranger versions 3.1.0 to 6.1.2 as data was generated. Samples from He, S et al., 2020 were re-processed from downloaded fastq files. Gene expression counts were corrected for ambient RNA using cellbender (Fleming *et al*., 2023) v0.2.0, using 0.6× and 2.5× cellranger identified cell count as –-expected-cells and –-total-droplets-included. Doublet removal was performed with scrublet (Wolock *et al*., 2019), cells with fewer than 500 umi, 300 genes, or more than 10% mitochondrial UMI were removed. Variable genes were identified using scanpy.pp.highly_variable genes flavour=seurat_v3, batch_key=project, span=0.5. Gene counts were normalized to 1e4 per cell and log transformed. 20 PCs were used for bbknn(Polański *et al*., 2020) batch correction. Corrected nearest-neighbors graph were used in UMAP projection and leiden clustering.

### Cell annotation

Cells were annotated with various references as input for manual curation: cello (Bernstein *et al*., 2021), Azimuth PBMC, Azimuth BBMC, Azimuth Motor cortex (Hao *et al*., 2021), celltypist Immune_All_High, Immune_All_Low (Domínguez Conde *et al*., 2022). Leiden clustering with high resolution plus manual annotation to merge clusters annotated to the same broad cell types or with few differential genes was used to assign cells to Lv1 annotations. After annotation of Lv1 cell types, each was sub-clustered to assign cells to Lv2 cell types following the same procedure as for the whole atlas Lv1 annotation with the difference of applying harmony batch correction.

### Meta-cells

Meta-cells were created within each Lv2 cell type using SEACells (Persad *et al*., 2023) v0.2.0 creating sqrt(n_cells) *2 meta-cells. UMI within genes or tCRE were summed from cells for each meta-cell and re-log-normalized.

### Annotation and quantification of tCREs in the atlas

To identify tCRE in the atlas data, the SCAFE *v1.0.0* (Moody *et al*., 2022) pipeline was applied to define and annotate tCREs. Briefly, for each library, the single nucleotide resolution 5’Cap TSS (CTSS) signals, including the number of UMI with unencoded Gs, were extracted from the alignment bam files generated from *cellranger*. The CTSS signals for all libraries of each “project” (as listed in Supplementary Table 1) were aggregated and used to define a set of TSS clusters for each project. For each project, the pooled CTSS signals were clustered using *paraclu* within SCAFE (Moody et al., 2022) using default parameters, with a cutoff set to ≥3 UMI of encoded-G supported TSS per TSS cluster. TSS clusters that are potentially strand invasion artifacts were removed (Moody et al., 2022). The remaining TSS clusters were further filtered using a logistic regression classifier trained with matched sc-5’-RNASeq and ATAC-Seq data implemented in SCAFE (Moody et al., 2022) at the logistic probability cutoff of 0.9. These remaining TSS clusters from each project, multiple hard filters were applied to remove the potentially artifactual clusters on the sense strand of the intronic and exonic regions of annotated genes, with ≥5 UMI within the cluster and ≥3 UMI at TSS cluster summit. A slightly more stringent cutoff was applied to the single nuclei libraries from project HCAJ0029 Brain tissues, with ≥10 UMI within the cluster, ≥5 UMI at TSS cluster summit and ≥5 UMI of encoded-G supported CTSS. These sets of filtered TSS clusters from all projects were merged using *bedtools merge* in a strand specific manner. The merged TSS clusters located within ±500nt of gene TSS annotated in GENCODEv32 were classified as proximal, or as distal otherwise. All TSS clusters were then extended 400nt upstream and 100nt downstream. These extended ranges were merged using *bedtools*, in a strand-specific manner for proximal TSS clusters and non-strand-specific manner for distal TSS clusters, as proximal-tCRE (P-tCREs) and distal tCREs (D-tCREs) respectively. The P-tCREs with its CTSS summit within 500nt of annotated gene TSS on the same strand would be annotated as promoter P-tCREs, and otherwise as flanking P-tCREs. It is noted that most flanking P-tCREs are on the opposite strand of the promoters, resembling promoter upstream antisense transcripts. For the D-tCREs that are located within the introns or exons of annotated genes, it will be “rescued” as promoter P-tCREs if, 1) its expression levels (number of UMIs within its TSS clusters) ≥5% of the expression levels of the corresponding gene (total number of UMI of all annotated promoter P-tCREs of the gene) and 2) ≥75% of its UMIs are on the same strand of the corresponding gene. In total, 8,791 D-tCRE were rescued as promoter P-tCREs, which can be considered as novel alternative promoters that are not annotated in GENCODEv32. In total, the above process yielded 81,829 P-tCREs and 96,400 D-tCRE, with 54,149 of 81,829 P-tCREs annotated as promoters. The average size of P-tCREs and D-tCREs are 771.12 nt and 608.01 nt, respectively. Expression of tCREs is quantified by counting the number of CTSS UMIs overlap with its constituent TSS clusters on the same stand.

### Defining of distal loci and HIFI loci

Distal loci is defined as a stretch of closely situated D-tCRE with a distance limit and P-tCREs were excluded from this analysis. To estimate an optimal distance, the closest distance of a D-tCRE to another was plotted against the rank and the tangle line of the curve was used to identify a cutoff at 17,065 nt. D-tCREs within this cutoff were ‘stitched’ together and defined 34,120 distal loci, which ∼31.% of them (n=10,547) contains ≥ 3 D-tCREs. A metric, “spreadness”, which quantifies the extent of evenness of UMI distribution across the constituent D-tCREs, is calculated as ratio of (the fraction of the the total number of UMI in the loci contributed by the highest expressed D-tCRE in) to the (total number D-tCREs in the loci). A distal locus with spreadness ≥ 4 is defined as evenly spread. The expression level of a distal locus is defined as the sum of the expression level of their constituent D-tCREs. To identify high intensity distal loci in each Lv1 cell type, the expression levels (log-normalized values) of each active distal loci (UMI count ≥ 1) were plotted against their ranks and the tangle line of the curve was used to identify a cutoff in each Lv 1 cell type, with a median of 8.36 among Lv1 cell types (Fig. 3i). Frequently interacting distal loci are defined in a cell type agnostic manner as distal loci with total number linked promoters (from its constituent D-tCREs) passing a cutoff of 75 (Fig. 3j), determined the same way as high intensity distal loci but plotting the number of linked promoters instead of expression levels. A distal locus that is 1) evenly spread, 2) frequently interacting, and 3) high intensity in one of the Lv 1 cell types were defined a HIFI loci, yielding 1,229 HIFI loci in total. Cell type-unrestricted and –restricted HIFI loci were defined by *k-mean* clustering of their binary presence/absence among Lv1 cell types with n=2 (Fig. 3l).

### Gini index

Gini index of all tCREs and all distal loci were calculated from the respective expression matrices on Lv2 cell types (n=180), using the *gini()* function implemented in the ‘*ineq*’ R package.

### Inferring Promoter-to-D-tCRE interactions (PD-links)

PD-links were inferred by integrating public chromatin interaction datasets with our tCRE atlas, including 1) Hi-C from ENCODE (ENCODE Project Consortium et al., 2020) (n=172), 2) H3K27ac HiChIP from HiChIPdb ((Zeng et al., 2023), n=129), and 3) pcHi-C from 3DIV (Yang et al., 2018) (n=28). Together with 4) tCRE co-activity estimated from our atlas (Methods). For 1), 2) and 3), the significant (FDR < 0.05) loops (at various resolutions) were taken as provided by the original sources. The details of the used chromatin interaction datasets were listed in Supplementary Table 12. For 1) and 3), the provided significant loops are at mixed resolutions, with mean of 5157.99 bp and 10739.5bp in 1) and 3) respectively. For 2), interactions at 5,000bp were chosen for our analyses. For 4), we estimated the co-activity of all tCRE pairs among all meta-cells across the whole atlas as well as the meta-cells within each Lv1 cell type, using *Cicero v1*.*3*.*4*.*11,* with the expression matrix of tCREs as input and ran in a non-binarized manner. For each pair of tCREs, the highest co-activity score among the above-mentioned scope was taken as the representative. A pair of promoter and D-tCRE is inferred as linked if both tCREs overlap a significant loop in 1), 2) or 3), or having a representative co-activity score ≥ 0.2. This analysis yields 466,079 linked promoter-D-tCRE pairs, involving 40,626 promoters with a median of 9 D-tCREs linked.

### Defining tCRE modules

tCRE modules are defined using cNMF (Kotliar *et al*., 2019) using the prepare, factorize, combine, consensus workflow for meta-cell tCRE expression. We used values of k from 50 to 250 in increments of 10, examining the stability/error plots to maximize the stability and number of components, selecting k=150 to define 150 modules providing tCRE spectra scores quantifying the contribution of each tCRE to the module.

### Processing of GWAS summary statistics

All GWAS summary statistics (n=63 traits and diseases) are listed in Supplementary Table 7. Briefly, GWAS summary statistics were obtained from (1) UK biobank heritability browser (https://nealelab.github.io/UKBB_ldsc/index.html), (2) Dr. Alkes Price group site (https://alkesgroup.broadinstitute.org/) and (3) other sources (refer to Supplementary Table 7). Summary statistics obtained from (1) and (2) were directly used for heritability enrichment analyses, while the summary statistics obtained from (3) were pre-processed using “*munge_sumstats*.*py”* scripts in *LDSC* software.

### Trait heritability enrichment in CREs

For analysis in Fig. 1i, Fig. 3n, Fig. 3o left column, Supplementary Fig. 9 and Supplementary Fig. 10, enrichment of trait heritability in CREs was assessed by stratified LD score regression (S-LDSC) implemented in *LDSC* software. Annotation files and LD score files were generated for each set of CREs using the “*make_annot*.*py*” and “*ldsc*.*py*” scripts using default parameters. Each set of CREs was added onto the 97 annotations of the baseline-LD model v2.2 and heritability enrichment (i.e., ratio of proportion of heritability to proportion of SNP) for each trait was estimated using the “*ldsc*.*py*” script with *“--h2”* flag in default parameters. For analysis in Fig. 3o, middle and right column, as well as the heritability enrichment in modules (described below), which involve the comparison of relative heritability enrichment between two sets of CREs, we used the “specifically expressed genes” approach (LDSC-SEG) implemented in *LDSC* software. Briefly, two sets of tCREs, one defined as “foreground” e.g. HIFI loci, was compared against a “background” tCRE set, e.g. non-HIFI loci. Annotation files and LD score files were generated for each set of “foreground” and “background” tCREs using the “*make_annot*.*py*” and “*ldsc*.*py*” scripts using default parameters. These foreground and background annotations were added onto the 53 annotations of baseline-LD model v1.2 and the contribution of “foreground” tCREs to trait heritability (i.e., regression coefficient) for each trait was estimated using the “*ldsc*.*py*” script with *“--h2-cts”* flag in default parameters.

### Trait heritability enrichment in modules

The extent of heritability enrichment for each trait in each module was quantified using the LDSC-SEG approach similar to the approach mentioned above, with the 53 annotations baseline-LD model v1.2. Briefly, for each module, the top 15000 tCREs ranked by the module contribution score (i.e. spectra) derived from cNMF was used as the ‘foreground’ and the rest of the tCREs were used as the ‘background’. The ‘‘foreground’ tCRE regions were compared against the ‘background’ tCRE regions by running *ldsc*.*py –-h2-cts,* yielding a p-value and a regression coefficient for each trait-module pair. The value of –log10(P) was as a score which is then further trimmed, scaled and powered within each trait as follows: 1) score of the modules with regression coefficient < 0 or p-value > 0.1 were set to zero; 2) the trimmed score was raised to the power of 1.5 to increase the contrast of high and low levels of heritability enrichment; 3) the powered score was scaled to the maximum score within the trait. This yields a value of 0 to 1 within each trait across all modules, which was then used as the weight to calculate the weighted sum of module usage for ICE-CREAM score described below.

### ICE-CREAM score

In essence, the ICE-CREAM (**<u>I</u>**ndividual **<u>C</u>**ell **<u>E</u>**nrichment of **<u>CRE</u> <u>A</u>**ctivity **<u>M</u>**odule) score, for a particular trait in a particular single-cell (or meta-cell) was calculated as the sum of module usage, each weighted by the extent of trait heritability enrichment in the corresponding module. Briefly, module usages were calculated for meta-cells (or single-cells) by running *cnmf_obj.refit_usage*(expr, spectra). The usage for each module in a cell is then weighted (i.e. multiplied) by the extent of heritability enrichment (explained above) for the corresponding module for a given trait. The weighted sum of all modules thus yields a score for each trait in each cell. To quantify the statistical significance of this score, a null distribution of the score is generated by permutation of the module usage. Briefly, the tCRE expression values were shuffled within 5 expression bins 1000 times to generate 1000 expression levels-matched random expression matrices as the input for rerunning *cnmf_obj.refit_usage,* yielding 1000 permuted module usage matrices. Weighted sums were then recalculated for 1000 times, while keeping the extent of heritability enrichment fixed, yielding a null distribution of the score. The observed score for a given trait in a cell was then compared against the corresponding null distribution, yielding a Z-score and thus a one-tailed p-value (P), using *scipy.stats.norm.sf*. The ICE-CREAM score is then calculated as –log10(P) yielding a non-negative value.

### Differential gene/tCRE expression and differential tCRE usage

Differential expression for gene or tCRE at Lv1 and Lv2 cell types were performed with *scanpy.tl.rank_genes_groups* with method=‘*t-test’*. In promoter usage analysis, promoters were considered if they had more than 10 UMI and >=5% of the UMI when summing all promoters assigned to a gene. The proportion of UMI from each promoter was calculated per meta-cell to give a promoter usage score. This score was visualized and used as input for differential expression testing to assign cell type enriched usage.Similarly to promoter usage, D-tCRE usage was calculated for each gene, using all D-tCRE that were linked with a gene promoter.

### Motif analysis

ChromVAR (Schep *et al*., 2017) using JASPAR2018 (Khan *et al*., 2018) motifs was applied to the tCRE meta-cell based matrix to estimate motif activity in each meta-cell. The motifbreakR R package (Coetzee *et al*., 2015) was used to assess the severity of SNP disruption of JASPAR2018 TFBMs.

### Gene Set Enrichment Analysis (GSEA)

fgsea v1.28 (Korotkevich *et al*., 2021) was used to score enrichment of gene sets from MSigDB (Hallmarks, Reactome, KEGG, GO biological processes and molecular functions) using maximum cNMF ‘gene spectra’ scores from promoters assigned to genes to rank genes.

### Cell Set Enrichment Analysis (CSEA)

fgsea was applied to groups of meta-cells within the atlas ranked by trait scores at two levels: Lv1 cell types within the ranking of the whole atlas, or Lv2 cell types within subsets of the atlas (stromal, immune, neural, epithelial).

### Defining trait associated SNPs, tCREs and genes with functional contexts

To define trait-associated SNPs, genome-wide significant lead variants (p < 5×10^−8^) were extracted from the 63 summary statistics listed in Supplementary Table 7. To increase coverage, additional genome-wide significant lead SNPs for each trait (by matching of ontology terms listed in Supplementary Table 7) were also extracted from extra GWAS studies from NHGRI-EBI GWAS Catalog (https://www.ebi.ac.uk/gwas/) (release r2023-06-03). The SNPs within the LD block of the GWAS lead SNPs (i.e., proxy SNPs) were searched for using *PLINK v1*.*9* with an r^2^ ≥0.2 within ±500kb in matched population panels of Phase 3 1000 Genomes Project downloaded from MAGMA website (http://ctg.cncr.nl/software/MAGMA/ref_data/). These lead and proxy SNPs are referred to as trait associated SNPs. Trait associated SNPs residing in a tCRE are then linked to a gene if the tCRE is the gene promoter or is a D-tCRE linked to genes through the mentioned P-D links. SNPs, and tCRE are further filtered to be enriched within trait relevant cell types – significant in Lv1 cell type CSEA to select relevant cell types, tCRE defined as enriched in cell type by significant Wilcoxon test or Pearson’s correlation with trait ICE-CREAM score > 0.5 across the whole atlas. Log normalized tCRE expression, gene expression, and distal loci expression are correlated with ICE-CREAM scores across meta-cells. Filtering for SNPs within relevant motifs: SNPs scored as disrupting TFBM by motifbreakR are listed as motif disrupting if the motif activity score across the atlas by chromVAR are significantly enriched in the same trait relevant cell types.

## Data availability

Data used in the initial cell line and PBMC comparisons are available in the ArrayExpress database (http://www.ebi.ac.uk/arrayexpress) under accession numbers: E-MTAB-10385 (sc-end5-dT, sc-end5-rand and sc-end3-dT for DMFB, iPSC), E-MTAB-10378 (sc-end5-dT for PBMC), E-MTAB-10381 (bulk-ATAC-seq for DMFB, iPSC), E-MTAB-10382 (sc-ATAC-seq for PBMC), E-MTAB-10383 (bulk-RNA-seq for DMFB, iPSC), E-MTAB-10384 (bulk-CAGE for DMFB, iPSC).

A genome browser view for the tCRE atlas are available at: https://jon-bioinfo.github.io/TCRE_Atlas/igv.html

Supplementary figures and tables are available at: https://doi.org/10.6084/m9.figshare.c.6926944

A cellxgene web portal, the processed data and the codes for data analyses will be made available for upon publication of the manuscript in a journal.

Due to patient data confidentiality sequencing data from the Single Cell Medical Network in Japan are not provided.

## Acknowledgements

This publication is part of the Human Cell Atlas (www.humancellatlas.org/publications) and the Single Cell Medical Network of Japan. This research was supported by a research grant to the RIKEN Center for Integrative Medical Sciences (IMS) from the Ministry of Education, Culture, Sports, Science and Technology (MEXT). We would like to extend our thanks to Chitose Takahashi, Nozomi Moritsugu, Hiroko Kinoshita, Tsugumi Kawashima from RIKEN IMS for assistance in single-cell RNA sequencing, to Teruaki Kitakura and Nobuyuki Takeda from RIKEN IMS for their contribution in the information infrastructure management for this project, and to Shiho Nakamura, Fumiko Ozawa, Mitsutoshi Tano for technical supports. We further acknowledge the Japan Science and Technology Agency (CREST-JPMJCR2011 to Taishin Akiyama; Forrest-21457195 to Tomohisa Sujino; JPMJIH1504 to Hiroshi Kawasaki), Grants-in-Aid from the Japanese Society for the Promotion of Science (JSPS) (21K18272 and 23H02899 to Tomohisa Sujino; 22K15736 and 21H05278 to Satoru Morimoto; 21H02853 to Ken-ichiro Kubo; 22K15203 to Satoshi Yoshinaga), Japan Agency for Medical Research and Development (AMED) (JP22ek0410079 and JP19ek0410046 to Hiroshi Kawasaki; JP22ek0109616, JP23ek0109651, JP23ek0109648, JP23kk0305024, JP23bm1423020, JP23bm1123046 and JP23bm1423002 to Satoru Morimoto and Hideyuki Okano; JP21wm0425019 to Masaki Takao). National Center of Neurology and Psychiatry (NCNP) biobank is partly supported by a grant from AMED (GAPFREE4-JP21ak0101151) and Intramural Research Grant (3-1) for Neurological and Psychiatric Disorders of NCNP.

## Author contributions

JW.S, CC.H. (Coordination, manuscript writing, study design, analysis), J.M. (Manuscript writing, data analysis), Y.A., P.C. (Coordination, study design), JC.C., J.L., C.T., CW.Y. (Data analysis), A.H., Mi.T., Ta.K. (Data management and coordination), Tu.K., M.K., I.K., T.H., S.N., Ko.O., F. LR., Y.S. (Performed experiments), T.A., N.A., M.A., A.FN., Mi.H., K.H., Mi.H., Y.I., K.I., H.K., Tos.K., Tom.K., K.K., Y.K., R.M., T.M., S.M., A.N., J.N., Hi.O., Ya.O., N.S., H.S., K.S., T.S., A.S., H.T., M.Taka, M.Take, T.T., K.Y., S.Y. (Sample procurement)

## Declaration of interests

The authors declare no competing interests.

## Notes

https://doi.org/10.6084/m9.figshare.c.6926944

### Competing Interest Statement

The authors have declared no competing interest.

